# The functional connectivity of the human claustrum according to the Human Connectome Project database

**DOI:** 10.1101/705350

**Authors:** Lluviana Rodríguez-Vidal, Sarael Alcauter, Fernando A Barrios

## Abstract

The claustrum is an irregular and fine sheet of grey matter in the basolateral telencephalon present in almost all mammals. The claustrum has been the object of several studies using animal models and more recently in human beings using neuroimaging. Believed to be involved in cognition and disease such as fear recognition, suppression of natural urges, multisensory integration, conceptual integration, seizures, multiple sclerosis, and Parkinson’s disease. Nevertheless, the function of the claustrum still remains unclear. We present a functional connectivity study of the claustrum in order to identify its main networks. Resting state functional and anatomical MRI data from 100 healthy subjects were analyzed; taken from the Human Connectome Project (HCP, NIH Blueprint: The Human Connectome Project), with 2×2×2 *mm*^3^ voxel resolution. Positive functional connectivity was found (p<0.05, FDR corrected) between the claustrum and the insula, anterior cingulate cortex, pre-central and postcentral gyrus, superior temporal gyrus, and subcortical areas. Our findings coincide with the results previously reported in both animal models and studies with humans. Showing the claustrum as a well-connected structure not only structurally but also functionally. Evidencing the claustrum as a node participating in different neural networks.

## Introduction

The claustrum is an irregular and fine sheet of grey matter in the basolateral telencephalon present in almost all mammals. The claustrum is separated from the insular cortex by the extreme capsule and medially from the lentiform nucleus by the external capsule^1–7^. Figure 1 shows in an axial view the anatomical location of the human claustrum. Research in humans using diffusion tensor imaging (DTI) have revealed cortical connections with the claustrum, which possesses projections to A) prefrontal cortex, BA 8, 9, 10, 11, 12 and 34; B) visual cortex, BA 17, 18, 19 and 39; C) sensoriomotor cortex, BA 7, 5, 1/2/3, 4, 6 and 8; and D) language areas BA 44, 45 and 31; as well as with orbitofrontal cortex, temporal cortex, basal ganglia and amygdala^3,7,8^ using DTI in 100 healthy subjects, Torgerson et al.^8^ found that the claustrum has the highest connectivity in the brain by regional volume. The literature about the claustrum – including studies in animal model and in humans – has evidenced the vast anatomical connections between the claustrum and the entire cerebral cortex as well as with the subcortical structures^1–4,6–9^.

**Figure 1.**
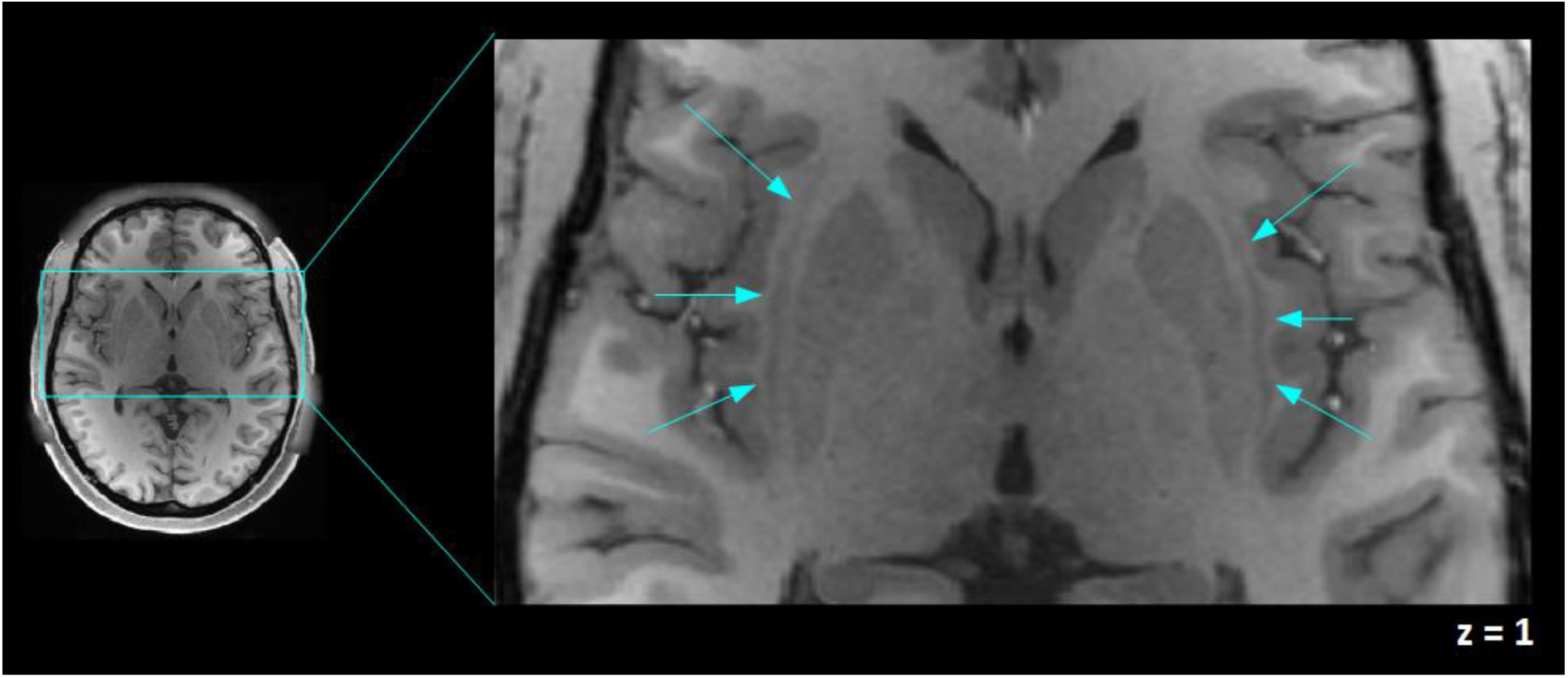
An axial view of the human claustrum. The human claustra are shown (signaled by blue arrows) between the external and extreme capsule, left and right claustra in a T1-weighted image of one of the participants are shown in MNI coordinates. This figure was acquired using Fslview, from the FMRIB software Library (FSL) tools v5.0^35^.

Based on its cellular composition and wide structural connectivity, the claustrum has been described as a “cornerstone of sensory integration”^1^. In a similar way,^2^ proposed the claustrum as a crucial component integrating motor and sensory information coming from different modalities to assemble them in a single experience; in an analogy used by Crick and Koch, the claustrum is as an orchestra conductor, “[w]ithout the conductor, the players can still play but they fall increasingly out of synchrony with each other.”^2^; actually, this proposal has been one of the most influential to the neuroscience community ^3,7,13,29,30,31^. In addition, the claustrum has been proposed as a relevant structure in the segregation of attention^4,6^.

The claustrum have been involved in cognitive process such as fear recognition, suppression of natural urges, multisensory integration and conceptual integration. The involvement of the claustrum in such processes is the result of research questions directed to the cognitive processes previously mentioned more than the intrinsic question about the function of the claustrum by itself^10–13^. Although it is rarely reported in the medical literature the selective lesion of the claustrum; the claustrum has been involved in disease such as seizures, multiple sclerosis, Parkinson’s diseases, disruption of consciousness, auditory hallucinations and delusions^14–17^.

Resting-state functional connectivity (RSFC) has proved to be a useful tool to characterize the functional communication of specific brain areas. Examining resting state functional connectivity allows us to determine the correlation between the spontaneous activity of brain areas that are anatomically separated, observing if there is functional communication between brain regions, specifically, is defined as the temporal dependency of neurophysiological events of anatomically separated brain regions^18^. A recent study in humans using functional magnetic resonance imaging (fMRI) shown the functional connectivity between the claustrum and anterior cingulate cortex, prefrontal, visual and parietal cortices^19^. They also reported functional connectivity between the claustrum and posterior cingulate cortex, precuneus, angular gyrus, cuneus, visual cortex, and sensorimotor cortex, as well as with subcortical structures such as thalamus and nucleus accumbens among others, according to this study, their results suggest the association of the claustrum with cognitive control. Nevertheless, in order to have a clear or complete understanding on the function of the claustrum beyond of the theoretical proposals^2,4^ we will require an accumulation of many studies to find certitude about our conclusions.

One of the most challenging issues to study the human claustrum is its intricate anatomical location and its irregular form. Actually, the claustrum dimensions have been reported through a postmortem 3D reconstruction imaging study by Kapakin^5^, having approximately 35.5710 mm x 1.0912 mm x 16.0000 mm and a volume of 828.8346 mm^3^ for the right claustrum and 32.9558 mm x 0.8321 mm x 19.0000 mm with a volume of 705.8160 mm^3^ for the left claustrum. To this respect, Milardi et al.^7^, have reported similar values for the claustrum mean volume, 813.6 mm^3^ for the right claustrum (range 744 – 864) and 804.0 mm^3^ for the left claustrum (range 752 – 912).] We propose to approach these challenges by assessing the whole brain resting state functional connectivity of the claustrum using the WU-Minn Human Connectome Project (HCP) dataset. The functional datasets were acquired with a spatial resolution of 2 mm isotropic and a temporal resolution of 720ms at 3T^20–24^. In this study, we aimed to explore, by a seed driven analysis, the resting-state functional connectivity of the human claustrum based on a large cohort of healthy subjects.

## Materials and methods

### Subjects

The study included data from 100 healthy young adults with ages between 22 and 35 years old, representing healthy subjects who are expected to pass the age of major neurodevelopmental changes and have not onset the age of degenerative changes^22^. Subjects were selected from the 1200 adult subject that is part of the WU-Minn Consortium (Principal Investigators: David Van Essen and Kamil Ugurbil; 1U54MH091657) funded by the 16 NIH Institutes and Centers that support the NIH Blueprint for Neuroscience Research; and by the McDonnell Center for Systems Neuroscience at Washington University, WU-Minn Human Connectome Project (HCP NIH Blueprint: The Human Connectome Project, https://www.humanconnectome.org/study/hcp-young-adult/data-releases)^22^. HCP excluded subjects having history of psychiatric, neurological or neurodevelopmental disorder or substance abuse. A full and detailed description of the recruitment criteria is provided by Van Essen et al.,^22,23^. Our sample from the HCP data was restricted to 100 unrelated subjects with complete resting-state functional magnetic resonance imaging (rfMRI) and anatomical MRI datasets. The WU-Minn HCP Consortium obtained the full informed consent from all participants following the Code of Ethics of the World Medical Association. All the protocols for data acquisition, distribution and use of the dataset also complied with the Code of Ethics of the World Medical Association^23^.

### Image acquisitions and preprocessing

The HCP acquired anatomical and resting-state functional MRI data at Washington University using a customized Siemens 3T “Connectome Skyra” with a standard 32-channel head coil. Structural dataset acquisitions included T1-weighted (T1w) and T2-weighted (T2w) images with 0.7 mm isotropic resolution, FOV = 224×224 mm, matrix = 320, 256 sagittal slices in a single slab TR = 2400 ms, TE = 2.14 ms, TI = 1000 ms, FA = 8°. Resting-state fMRI were acquired at 2 mm isotropic resolution, TR=720ms, TE = 33.1 ms, slice thickness of 2.0 mm, 72 slices. Uğurbil et al.,^24^ and Glasser et al.,^20^ provide a full and detailed description of the HCP acquisition protocols. We include in our study the HCP dataset preprocessed by the HCP which due its spatial and temporal resolutions and differing distortions must be processed differently from standard neuroimaging data in order to achieve optimal results^20,24^. Pipelines developed by HCP include spatial distortion correction, motion correction, spatial registration and normalization to MNI coordinates, highpass filtering (cutoff=2,000s) and denoising of rfMRI data^20,21^. In addition, we use as confound regressors BOLD signal from the white matter, cerebrospinal fluid (CSF) mask, realignment and scrubbing parameters and band-pass filtering (0.01 – 0.08 Hz)^33,34^. In order to minimize potential signal contamination from adjacent structures, the timeseries data from regions of interest (ROIs) were not smoothed. The linear regression and filtering were carried out using *conn toolbox* (V18b, Functional connectivity toolbox, NITRC)^25^, SPM 12 software^26^ and MATLAB R2018a (https://www.mathworks.com). The analysis was executed using the standard resting state pipeline included in *Conn* to simplified posterior reproducibility.

### Mask of claustrum and functional connectivity

The T1 anatomical volumes were used to manually delineate right and left claustrum masks for each subject, in an axial plane, in a superior-to-inferior manner to identify the body of the claustrum and edited in the coronal and sagittal plane for accuracy (Figure 2). We transformed them into the rfMRI space, spatially averaged the claustrum masks and established the threshold (0.1) of the mask to transform it into a binary mask. The transformation of the mask was a linear transformation, Tri-linear interpolation method and was carried out by the Linear Image Registration Tool (FLIRT). All these processes were carried on through the FMRIB software Library (FSL) tools v5.0^35^. The averaged claustrum masks constituted our seeds to perform a seed-based functional connectivity analysis (Figure 3).

**Figure 2.**
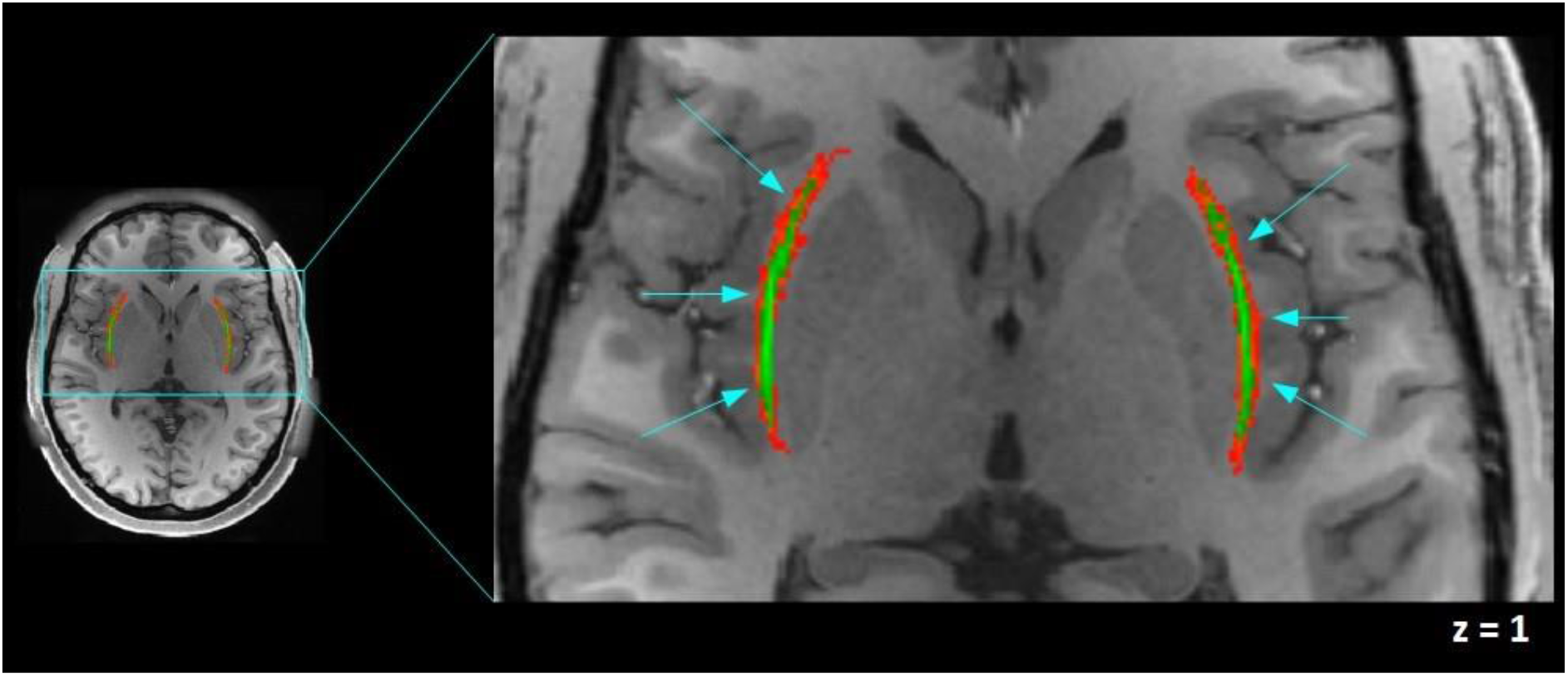
Axial view of the claustrum mask. The averaged mask of the claustrum is shown in green, left and right claustra mask on a T1-weighted image of one of the participants are shown in MNI coordinates. In green, our effective claustrum ROI, was estimated from the intersection of all subjects’ claustrum. In red, variability area due to subject differences. In our analysis we include only the green area. This figure was acquired using Fslview, from the FMRIB software Library (FSL) tools v5.0^35^.

**Figure 3.**
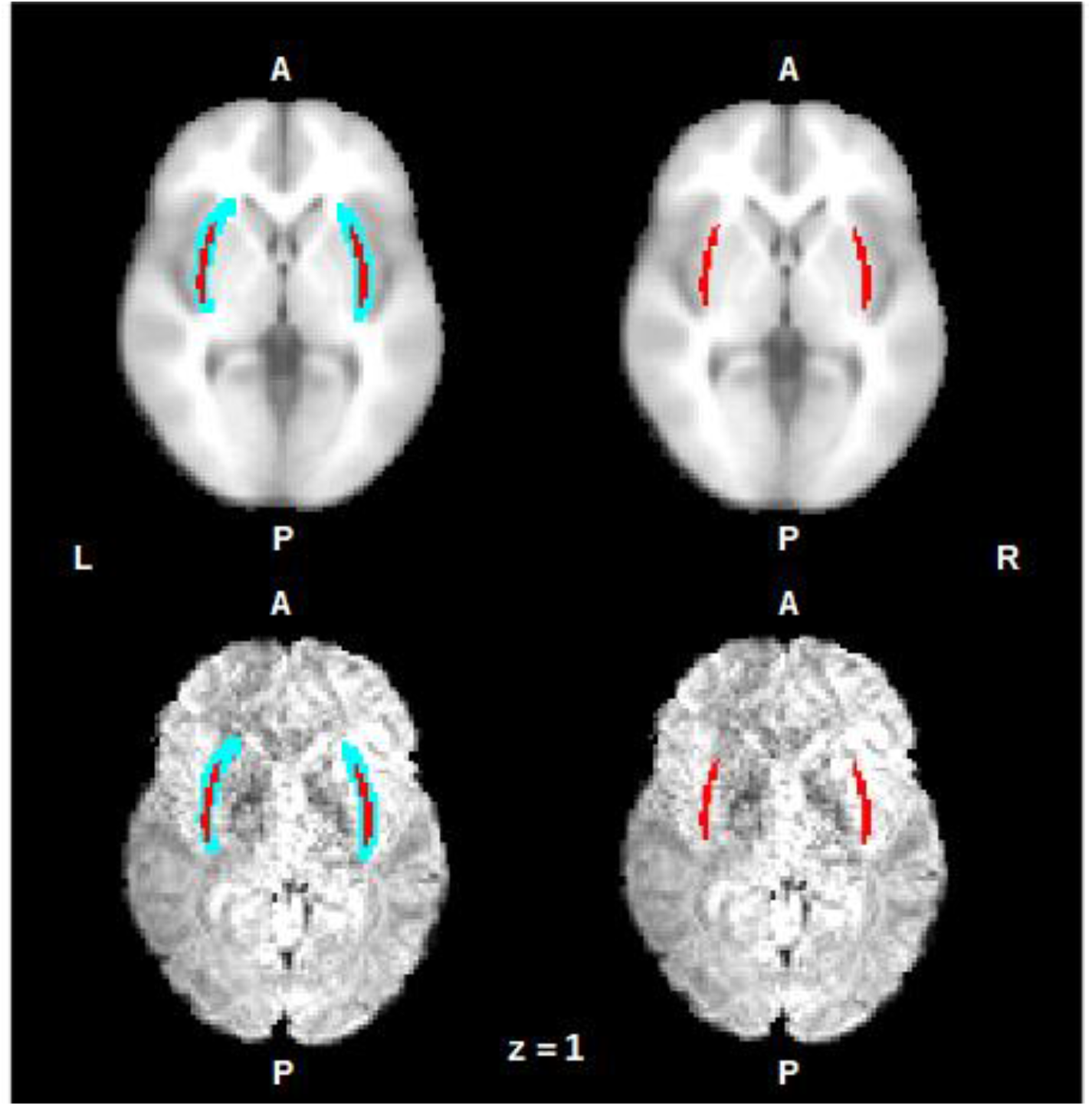
On an axial view the averaged mask of the claustrum is shown in red, left and right claustra mask on an anatomical MNI152 template (up) and on a representative subject functional image (down). The mask is shown after the transformation on the spatial functional resolution. On the left side: the blue area represents the variability area due to subject differences, on the right side, the red area represents the intersection of all subjects’ claustrum. R = right, L = left, A = anterior, P = posterior. This figure was acquired using Fslview, from the FMRIB software Library (FSL) tools v5.0^35^.

We performed a seed-to-ROI analysis using as seeds the right and left claustrum masks and 132 ROIs combining FSL Harvard-Oxford atlas cortical and subcortical areas and AAL atlas cerebellar areas, the analysis was carried on using Conn toolbox.

Functional connectivity maps were estimated using the *Conn toolbox* (V18b, Functional connectivity toolbox, NITRC)^25^, SPM 12 software^26^ and MATLAB R2018a. Pearson correlation coefficients were calculated within each subject in MNI space taking the seed time course and the time course of all other voxels (seed-to-voxel analysis) or the cortical/subcortical structures (seed-to-ROI analysis). The resultant correlation coefficient maps were converted to normally distributed scores using Fisher transform to permit a second-level analysis, in which, a one-sample t-test was performed. The resultant connectivity maps were corrected at map level p<0.05 using False Discovery Rate (FDR).

We carry out a comparative resting-state functional connectivity of the claustrum by contrasting functional connectivity of the left claustrum and right claustrum.

## Results

The claustrum was delimited by our averaged mask whose dimensions are 31mm x 1 mm x 17 mm. We closely coincide with the dimension described by Kapakin^5^; figures 2 y 3 show on an axial view the average mask used in the analysis, our effective claustrum ROI, was estimated from the intersection of all subjects’ claustrum.

In a seed-to-voxel analysis, we identified functional connectivity of the left claustrum (p<0.05 p-FDR corrected) with the following clusters: a) Left claustrum with a t–score 23.44, including left and right (l, r) precentral gyrus, postcentral gyrus (l, r), insular cortex (l, r), opercular cortex (l, r), anterior cingulate cortex, supramarginal gyrus (l, r), supplementary motor cortex (l, r), planum temporale (l, r), putamen (l, r), thalamus (l), amygdala (l). b) Left lingual gyrus (t–score 7.92), including intracalcarine cortex (l, r), precuneous cortex, cuneal cortex (r). c) Left occipital fusiform gyrus (t–score 6.79), including temporal occipital fusiform cortex (l). d) Left cuneal cortex (t–score 6.87). e) Left lateral occipital cortex (t–score 5.93) (Figure 4, Supplementary Table S1).

**Figure 4.**
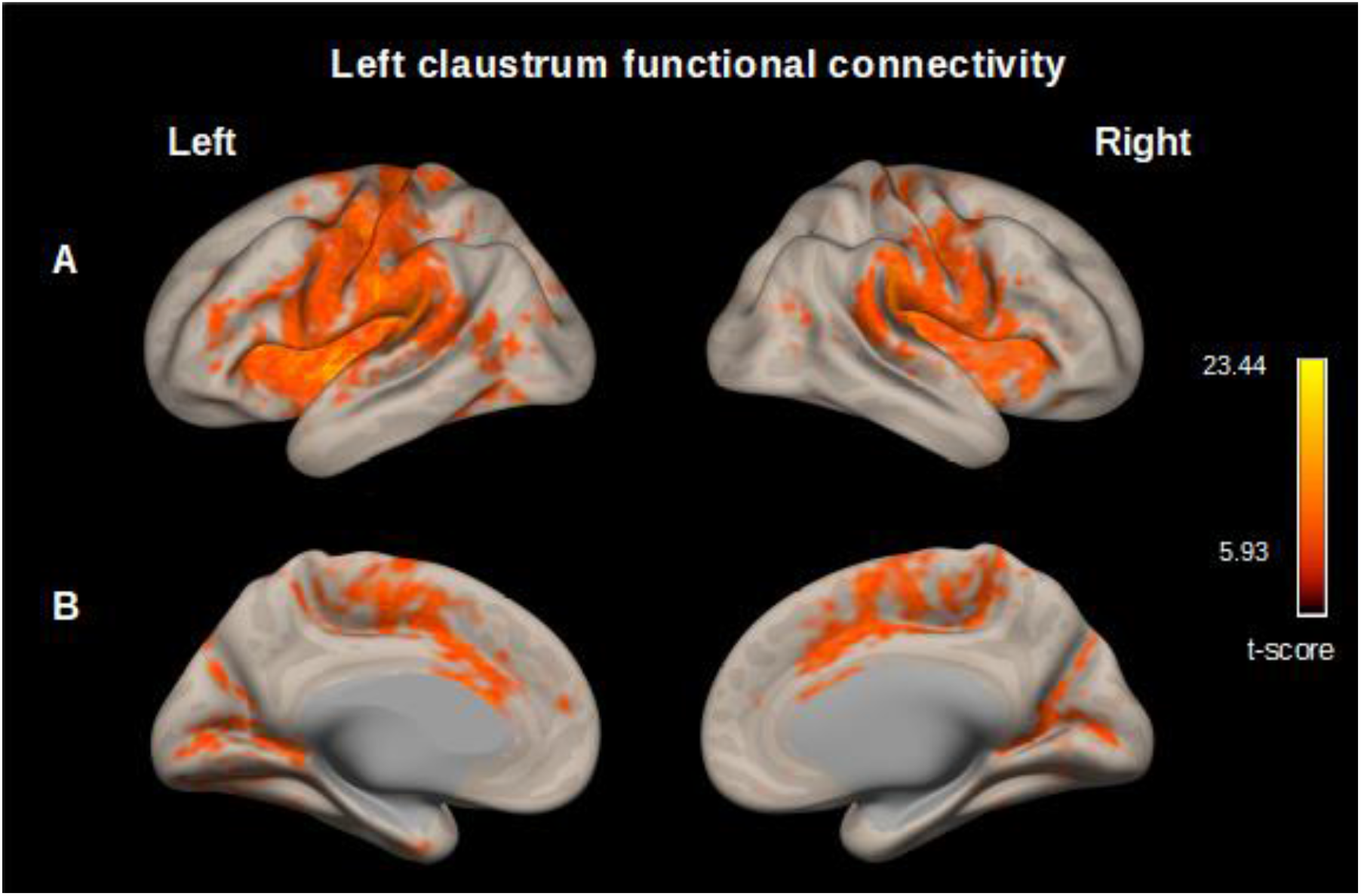
Map of the resting-state functional connectivity with left claustrum as seed. (p<0.05, p-FDR corrected) Row (A) Left, PreCentral Gyrus, postcentral Gyrus, insular cortex, opercular cortex supramarginal gyrus. Right, insular cortex, precentral and postcentral gyrus, supramarignal gyrus. (B) Left and right, cingulate gyrus (anterior division), occipital cortex. These functional connectivity maps were estimated using the *Conn toolbox* (V18b, Functional connectivity toolbox, NITRC)^25^.

We found functional connectivity between the right calustrum and the following clusters: a) Right claustrum (t–score 18.40) including precentral gyrus (r), postcentral gyrus (r), insular cortex (r), opercular cortex (r), supramarginal gyrus (r), anterior cingulate cortex, supplementary motor cortex (r), inferior frontal gyrus (r), planum temporale (r), putamen (r). b) Left planum polare (t–score 9.53) including opercular cortex (l), precentral gyrus (l), postcentral gyrus (l), insular cortex (l), planum temporale (l). c) Left precentral gyrus (t–score 7.17) including middle frontal gyrus (l). d) Right lingual gyrus (t–score 7.04), precuneous cortex, intracalcarine cortex (r), cuneal cortex (r), lingual gyrus (r) (Figure 5, Supplementary Table S2).

**Figure 5.**
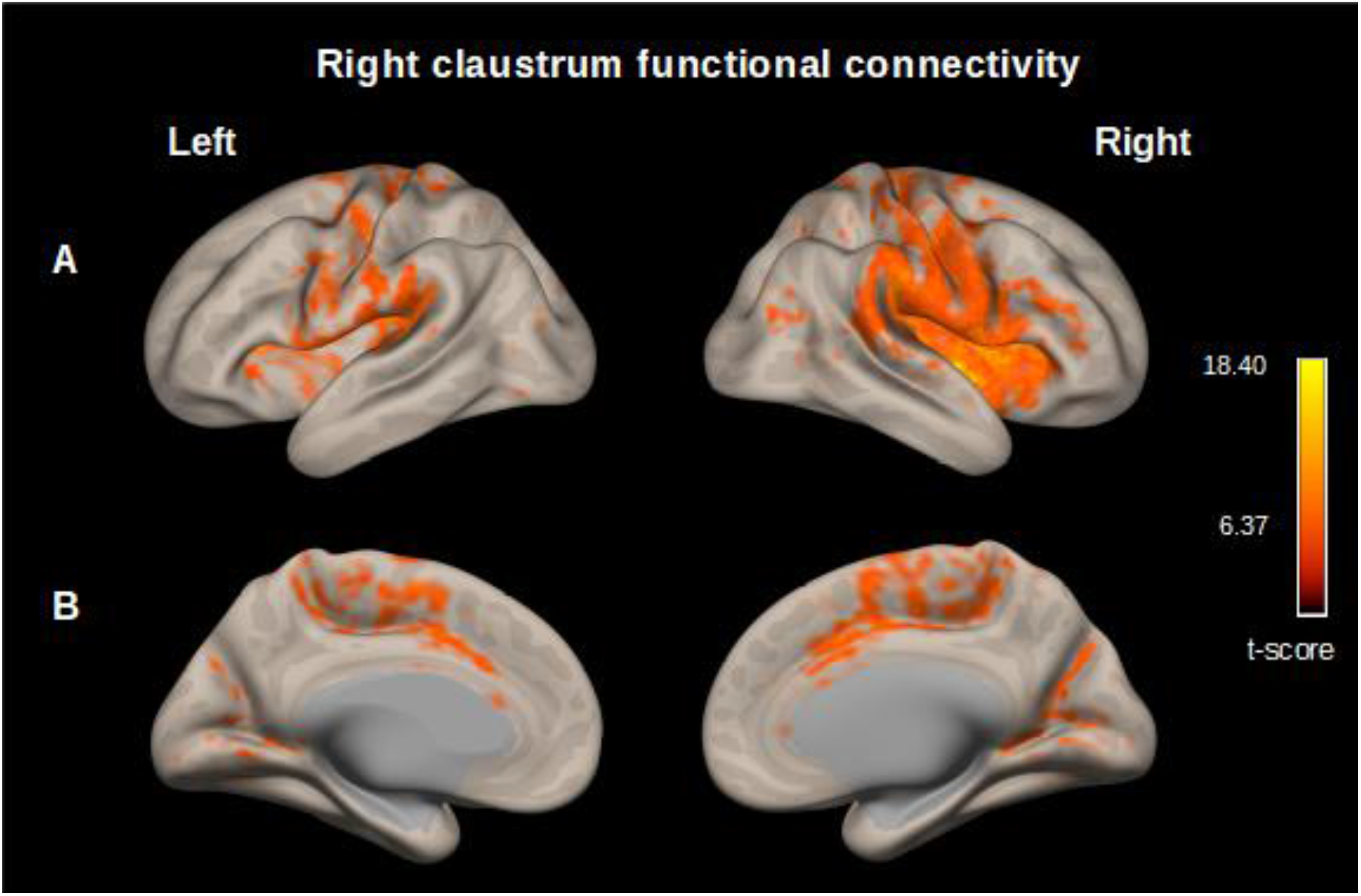
Map of the resting-state functional connectivity with right claustrum as seed. (p<0.05, p-FDR corrected) Row (A) Left, Planum polare, central opercular cortex, precentral gyrus, postcentral gyrus. Right, Precentral Gyrus, postcentral gyrus, insular cortex, central opercular cortex, supramarginal gyrus, parietal operculum cortex. (B) Left and right cingulate gyrus (anterior division) occipital cortex. These functional connectivity maps were estimated using the *Conn toolbox* (V18b, Functional connectivity toolbox, NITRC)^25^.

Left claustrum and right claustrum resting-state functional connectivity were contrasted in a seed-to-ROI analysis. When comparing left claustrum *versus* right claustrum functional connectivity, we obtained to the left claustrum cortical areas such as insular cortex, frontal orbital cortex, Hesch’l gyrus, opercular cortex, inferior frontal gyrus, frontal pole, paracingulate gyrus, temporal cortex, angular gyrus, planum temporale, middle frontal gyrus, temporal fusiform cortex left side in all cases, which imply an ipsilaterality. We found insular cortex, frontal pole and middle frontal right cortex to right claustrum, (for a complete description see Figures 6, and Supplementary Table S3) results were thresholded p<0.05 p-FDR corrected at seed-level.

**Figure 6.**
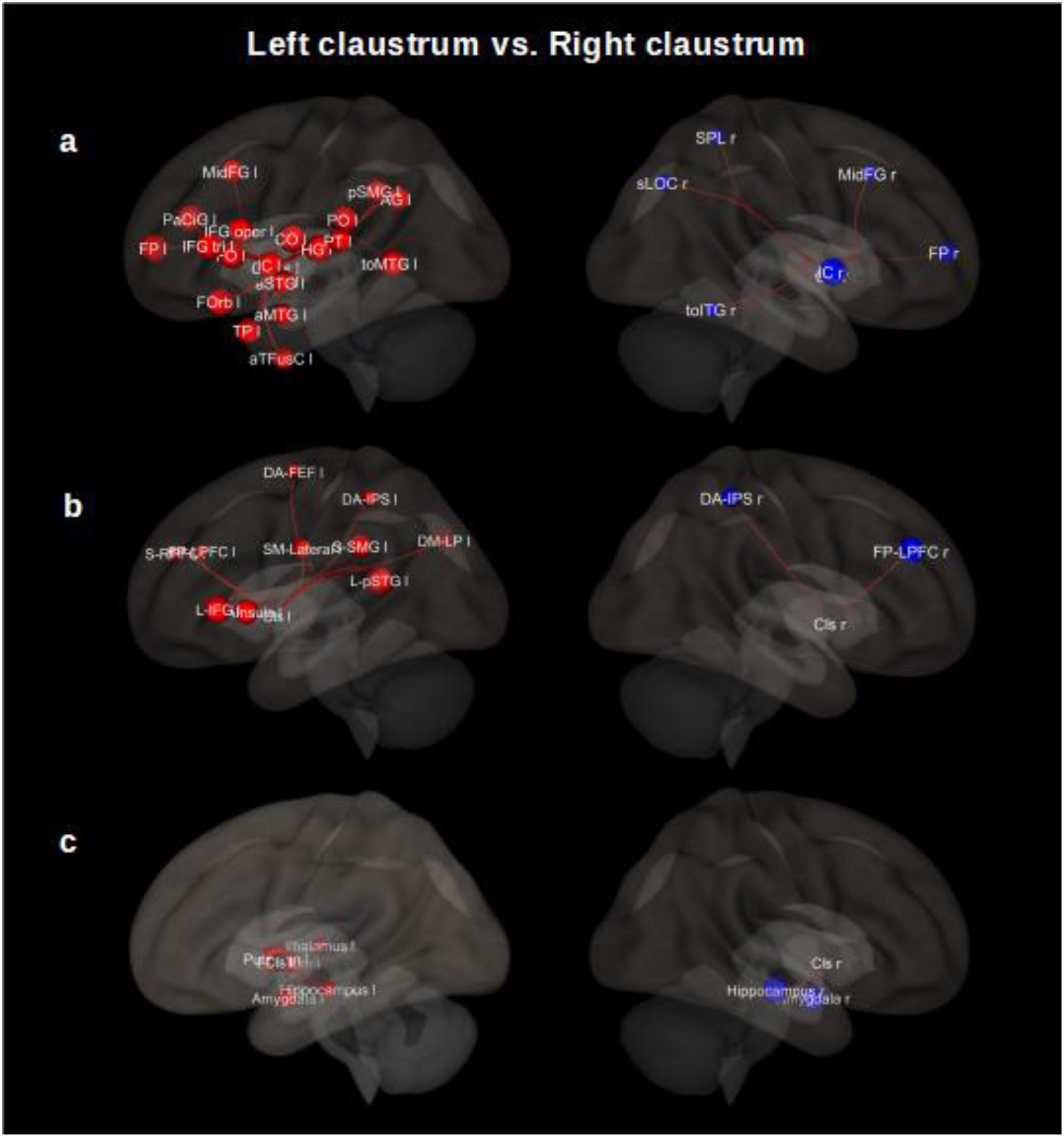
Comparison of the left claustrum with right claustrum resting-state functional connectivity. **Row (A)** shows cortical areas. Left (Cls – l > Cls – r), Insular cortex (IC l), frontal orbital cortex (FORrb l), Heschl’s gyrus (HG l), frontal operculum cortex (FO l), central opercular cortex (CO l), inferior frontal gyrus (IFG tri l), frontal pole (FP l), paracingulate gyrus (PaCiG l), middle temporal gyrus (toMTG l), parietal operculum cortex (PO l), inferior frontal gyrus (IFG oper l), temporal pole (TP l), angular gyrus (AG l), planum temporale (PT l), middle frontal gyrus (MidFG l), temporal fusiform cortex (aTFusC l), supramarginal gyrus (pSMG l), planum polare (PP l), middle temporal gyrus (aMTG l), superior temporal gyrus (aSTG l), superior frontal gyrus (SFG l), middle temporal gyrus (pMTG l), Precentral Gyrus (PreCG l), supramarginal gyrus (aSMG l), parahippocampal gyrus (aPaHC l), superior temporal gyrus (pSTG l), postcentral gyrus (PostCG l), parahippocampal gyrus (pPaHC l), temporal occipital fusiform cortex (TOFusC l), inferior temporal gyrus (pITG l), lateral occipital cortex (sLOC l), inferior temporal gyrus (aITG l), superior parietal lobule (SPL l). Right (Cls – r > Cls – l), insular cortex (IC r), frontal pole (FP r), lateral occipital cortex (sLOC r), middle frontal gyrus (MidFG r), superior parietal lobule (SPL r), inferior temporal gyrus (toITG r). **Row (B)** shows networks nodes. Left (Cls – l > Cls – r), languaje network (L): superior temporal gyrus (pSTG l), inferior frontal gyrus (IFG l), salience network (S): Insula (AI), supramarginal gyrus (SMG l), rostral prefrontal cortex (RPFC l), anterior cingulate cortex (ACC). Sensorio-motor lateral network (SM): lateral, dorsal attention network (DA): intraparietal sulcus (IPS l), frontal eye field (FEF L), Default mode network (DM): lateral parietal (LP). Fronto– Parietal network (FP): lateral prefrontal cortex (LPFC l). Right (Cls – r > Cls – l), Fronto–parietal network: lateral prefrontal cortex (LPFC r). Dorsal attention: Intraparietal sulcus (IPS r). The results are thresholded (p< 0.05, p-FDR corrected). Cls = claustrum, l = left, r = right, a = anterior division, p= posterior division, s = superior division, tri= pars triangularis, to = temporooccipital part, oper = pars opercularis. These 3D functional connectivity maps were proyected using the *Conn toolbox* (V18b, Functional connectivity toolbox, NITRC)^25^.

In our contrasting analysis, we also found functional connectivity with specific nodes of networks such as superior temporal gyrus and inferior frontal gyrus (language); anterior insular cortex, supramarginal gyrus and rostral prefrontal cortex (salience); sensoriomotor lateral; intraparietal sulcus and frontal eye field (dorsal attention); lateral parietal cortex (default mode); lateral prefrontal cortex (frontoparietal); for left claustrum and lateral prefrontal cortex (frontoparietal); Intraparietal sulcus (dorsal attention) for right claustrum. Considering subcortical regions, we obtained putamen, pallidum, amygdala, hippocampus and thalamus for left claustrum and hippocampus, amygdala for right one, (for a complete description see Figures 6, Supplementary Table S4 and Table S5) results were thresholded p<0.05 p-FDR corrected at seed-level.

## Discussion

Here, we carried out a seed-driven functional connectivity analysis in a sample of 100 healthy subjects. We found that the human claustrum is widely connected with cortical and subcortical brain areas. Left and right claustrum are mainly connected with precentral gyrus, postcentral gyrus, insular cortex, opercular cortex, supplementary motor area, anterior cingulate cortex, paracingulate cortex, frontal cortex, temporal and occipital cortex, putamen, hippocampus and amygdala. We found functional connectivity with specific nodes of the characterized brain networks such as salience, sensori-motor, language, dorsal attention, default mode, frontoparietal, which imply that the claustrum is not only well connected structurally but also functionally.

Our findings reveal that the claustrum maintains a functional connectivity with regions of the salience network, this could be justified not only due to the anatomical closeness of the claustrum with the insula, but also because they share ontogeny. Pirone *et al*.,^27^ using immunohistochemistry technique and samples of insular and temporal subunits of the human claustrum, revealed that the claustrum share ontogeny with the insular cortex, but not with putamen. Additionally, Remedios et al.,^28^ analyzed the neurons located in the auditory zone of the primate claustrum and arrived at the conclusion that the clausturm detects the occurrence of novel or salient stimulus. Despite of the fact that both the claustrum and the insula are related to salience, both differ in their functional connectivity patterns, when we contrasted both functional connectivity maps through a resting states analysis (Supplementary Fig. S1 and Fig. S2).

On the other hand, we found functional connectivity between the claustrum and sensori-motor areas, anterior cingulate cortex, frontal cortex, which is consistent with previous research that the claustrum participates in cognitive processes. Coactivation of the claustrum has been previously reported in different cognitive processes such as integration of sensory signal across the tactile and visual modalities, multisensory integration of conceptually related common objects, suppression of natural urges, retrieval fluency, cognitive control and task switching^10–13,19,29^. Claustral activation is reported in a fMRI study about the neuronal bases of the integration of auditory and visual stimuli, which, were conceptually related (congruent). The authors of this study concluded that the contribution of the claustrum was related to the congruence of stimuli, not only to the matching of these^13^.

To date, there are several hypotheses about the function of the claustrum; one of the most important in this regard was proposed by Crick and Koch^2^, who proposed the claustrum as an important hub which integrates stimuli from different sensory modalities. Hadjikhani et al.,^11^ researching neural bases of tactile–visual cross-modal transfer of shape in humans using neuroimaging, found that the claustrum has an important role in cross-modal matching. Their results suggest that the cross-modal adaptation is associated with multimodal converging brain areas such as anterior cingulate cortex, inferior parietal lobe, superior and mid temporal cortex, dorsolateral prefrontal cortex and insular cortex. Banati et al.,^10^ also found activation by tactile-visual stimuli in anterior cingulate cortex, dorsolateral prefrontal cortex, mid and superior temporal cortex and insula cortex. Our findings are consistent with these previous studies, anterior cingulate cortex, superior, temporal cortex and insular cortex are functionally connecting with the claustrum.

On the other hand, another important hypothesis about the function of the claustrum is related to segregation of attention^4,6^. The claustrum is considered as a hub for attention, according to this model, the prefrontal cortex modules to the claustrum in a top down process and the output of the claustrum modules the cortical representation of the sensory modalities. The claustrum controls which cortical output will be attended amplifying the preferred modality and reducing the other ones. This process would be modulated by additional subcortical inputs related to the context and emotion hippocampus and amygdala respectively. Our results are compatible with this model, since we found a positive correlation between the claustrum and the prefrontal cortex, anterior cortex cingulate, pre and postcentral gyrus, amygdala and hippocampus, which, according to the model of Goll et al.^4^ are implied in the segregation of attention^4^.

Also, our results show that correlation maps are partially lateralized; cortical, subcortical areas and nodes of salience network, language and dorsal attention display stronger and ipsilateral connection with left claustrum in comparison with right claustrum. Torgerson et al.^8^ in an DTI study of 100 neurotypical human subjects reported an asymmetry, right claustrum volume less than left claustrum volume. On the other hand, it was reported by Cascella et al.^15^ that severity of delusions in schizophrenia is correlated with the reduction in left claustral volume.

These results suggests that the claustrum maintains a wide functional connectivity with brain areas associated with cognitive process, such as temporal cortex pertaining to language network, lateral prefrontal cortex and intraparietal sulcus related to fronto-parietal and dorsal attention networks respectively; which suggests that the claustrum is participating not only in task switching^19^, but also in processed related to flexible modulation of other brain networks^36^ and process related to attention^37^.

This study contributes to the understanding of this thin but well connected structure: the claustrum. The use of the HCP datasets supplied us with important advantages to our study, particularly with its spatial and temporal resolution; and to the standardized preprocessing accomplished by the HCP group. Nevertheless, some limitations should be noted. The age range chosen (22-35 years-old) could only represent the resting state functional connectivity of the claustrum in healthy adults who are not experiencing the major neurodevelopmental changes nor neurodegenerative changes^22^; but leave out the functional connectivity claustrum in other age range. Our analyses are limited to the resting-state and do not include task-related functional imaging. Also, we focus on present the connectivity of the average claustrum, we did not consider handedness or gender related effects. We have carefully delineated a mask corresponding to the claustrum, nevertheless, this anatomical structure is itself fine, irregular and its location complicates a finer analysis. Technical constrains have limited the detailed study of the human claustrum *in vivo*; it is due to the anatomical location since it is located between two white matter structures (extreme and external capsules), irregular form taking the concavity form of the insular cortex and the convexity of the putamen, even this fine sheet-structure is not visible in some low-resoltution MR imaging^32^; we consider the advantages of achieve an fMRI study with human subjects *in vivo*, since most of the studies about the claustrum to the date are carried out in animal model.

In conclusion, the claustrum is a highly connected brain structure, hardly to study due its anatomical location and irregular form. Neuroimaging offers a powerful tool to explore its function non-invasively and *in vivo*, despite its limited resolution. Making use of open access datasets, we present an approximation of the resting-state functional connectivity of the claustrum, which maintains positive functional connectivity not only with cortical brain areas, but subcortical and different brain networks such as salience, sensorimotor, language, and attention.

## Acknowledgements

We thank the “Programa de Doctorado en Ciencias Biomédicas” of the “Universidad Nacional Autónoma de México (UNAM)” and the Consejo Nacional de Ciencia y Tecnología, (CONACYT) for the doctoral fellowship no. 477467 (Ll R-V) supported by the National Council of Science and Technology, Mexico (Consejo Nacional de Ciencia y Tecnología, CONACYT) and the PAPIIT program DGAPA-UNAM for the grant IN202119. Data were provided by the Human Connectome Project, WU-Minn Consortium (Principal Investigators: David Van Essen and Kamil Ugurbil; 1U54MH091657) funded by the 16 NIH Institutes and Centers that support the NIH Blueprint for Neuroscience Research; and by the McDonnell Center for Systems Neuroscience at Washington University. LlR-V is grateful to N. Aranda-López for her administrative support. We are also grateful to M.Sc. Leopoldo González-Santos and Dr. Erick H. Pasaye for technical support and Dr. M.C. Jeziorski for his revision of the manuscript.

## Author contributions

LlR-V, SA and FAB developed the study concept, contributed to the study design. Data analysis and interpretation was performed by LlR-V, SA and FAB. All work was done under the supervision of FAB. LlR-V wrote the manuscript draft. LlR-V prepared all the figures. All authors provided critical revisions and approved the final version of the manuscript.

## Data availability

All data related to this research is public and accessible at the Human Connectome Project site https://www.humanconnectome.org/study/hcp-young-adult/data-releases

## References

1. Edelstein, L. R. & Denaro, F. J. The claustrum: a historical review of its anatomy, physiology, cytochemistry and functional significance. Cell Mol Biology Noisy-le-grand France 50, 675–702 (2004).

2. Crick, F. C. & Koch, C. What is the function of the claustrum? Philosophical Transactions Royal Soc B Biological Sci 360, 1271–1279 (2005).

3. Fernández-Miranda, J. C., Rhoton, A. L., Kakizawa, Y., Choi, C. & Alvarez-Linera, J. The claustrum and its projection system in the human brain: a microsurgical and tractographic anatomical study. J Neurosurg 108, 764–774 (2008).

4. Goll, Y., Atlan, G. & Citri, A. Attention: the claustrum. Trends Neurosci 38, 486–495 (2015).

5. Kapakin, S. The claustrum: three-dimensional reconstruction, photorealistic imaging, and stereotactic approach. Folia morphologica 70, 228–234 (2011).

6. Mathur, B. N., Caprioli, R. M. & Deutch, A. Y. Proteomic analysis illuminates a novel structural definition of the claustrum and insula. Cereb Cortex 19, 2372–2379 (2009).

7. Milardi, D. et al. Cortical and subcortical connections of the human claustrum revealed in vivo by constrained spherical deconvolution tractography. Cereb Cortex 25, 406–414 (2015).

8. Torgerson, C. M., Irimia, A., Goh, S. Y. M. & Horn, J. D. V. The DTI connectivity of the human claustrum. Hum Brain Mapp 36, 827–838 (2015).

9. Qadir, H. et al. Structural Connectivity of the Anterior Cingulate Cortex, Claustrum, and the Anterior Insula of the Mouse. Front Neuroanat 12, 100 (2018).

10. Banati, R. B., Goerres, G. W., Tjoa, C., Aggleton, J. P. & Grasby, P. The functional anatomy of visual-tactile integration in man: a study using positron emission tomography. Neuropsychologia 38, 115–124 (2000).

11. Hadjikhani, N. & Roland, P. E. Cross-modal transfer of information between the tactile and the visual representations in the human brain: A positron emission tomographic study. J Neurosci 18, 1072–1084 (1998).

12. Lerner, A. et al. Involvement of insula and cingulate cortices in control and suppression of natural urges. Cereb Cortex 19, 218–223 (2009).

13. Naghavi, H. R., Eriksson, J., Larsson, A. & Nyberg, L. The claustrum/insula region integrates conceptually related sounds and pictures. Neurosci Lett 422, 77–80 (2007).

14. Kalaitzakis, M. E., Pearce, R. K. B. & Gentleman, S. M. Clinical correlates of pathology in the claustrum in Parkinson’s disease and dementia with Lewy bodies. Neurosci Lett 461, 12–15 (2009).

15. Cascella, N. G., Gerner, G. J., Fieldstone, S. C., Sawa, A. & Schretlen, D. J. The insula-claustrum region and delusions in schizophrenia. Schizophr Res 133, 77–81 (2011).

16. Cascella, N. G. & Sawa, A. The Claustrum. 237–243 (2014) doi:10.1016/b978-0-12-404566-8.00009-x.

17. Wada, J. A. & Kudo, T. Involvement of the claustrum in the convulsive evolution of temporal limbic seizure in feline amygdaloid kindling. Electroencephalography and clinical neurophysiology 103, 249–256 (1997).

18. Heuvel, M. P. van den & Pol, H. E. H. Exploring the brain network: A review on resting-state fMRI functional connectivity. Eur Neuropsychopharm 20, 519–534 (2010).

19. Krimmel, S. R. et al. Resting state functional connectivity and cognitive task-related activation of the human claustrum. Neuroimage 196, 59–67 (2019).

20. Glasser, M. F. et al. The minimal preprocessing pipelines for the Human Connectome Project. Neuroimage 80, 105–124 (2013).

21. Smith, S. M. et al. Resting-state fMRI in the Human Connectome Project. Neuroimage 80, 144–68 (2013).

22. Essen, D. C. V. et al. The Human Connectome Project: a data acquisition perspective. Neuroimage 62, 2222–2231 (2012).

23. Essen, D. C. V. et al. The WU-Minn Human Connectome Project: An overview. Neuroimage 80, 62–79 (2013).

24. Ugurbil, K. et al. Pushing spatial and temporal resolution for functional and diffusion MRI in the Human Connectome Project. Neuroimage 80, 80–104 (2013).

25. Whitfield-Gabrieli, S. & Nieto-Castañón, A. Conn: A Functional Connectivity Toolbox for Correlated and Anticorrelated Brain Networks. Brain Connectivity 2, 125–141 (2012).

26. Friston, K. J., Jezzard, P. & Turner, R. Analysis of functional MRI time-series. Hum Brain Mapp 1, 153–171 (1994).

27. Pirone, A. et al. Topography of Gng2-and NetrinG2-expression suggests an insular origin of the human claustrum. Plos One 7, e44745 (2012).

28. Remedios, R., Logothetis, N. K. & Kayser, C. A role of the claustrum in auditory scene analysis by reflecting sensory change. Frontiers Syst Neurosci 8, 44 (2014).

29. Volz, K. G., Schooler, L. J. & Cramon, D. Y. von. It just felt right: the neural correlates of the fluency heuristic. Conscious Cogn 19, 829–837 (2010).

30. Tian, F., Tu, S., Qiu, J., Lv, J. Y., Wei, D. T., Su, Y. H., and Zhang, Q. L. Neural correlates of mental preparation for successful insight problem solving. Behav. BrainRes. 216, 626–630 (2011).

31. Koubeissi, M.Z., Bartolomei, F., Beltagy, A., Picard, F., 2014. Electrical stimulation of a small brain area reversibly disrupts consciousness. Epilepsy Behav. 37, 32–35.

32. Torgerson, C.M., Van Horn, J.D., 2014. A case study in connectomics: the history, mapping, and connectivity of the claustrum. Front Neuroinform 8, 83. doi:10.3389/fninf.2014.00083

33. Biswal, B., Yetkin, F.Z., Haughton, V.M., Hyde, J.S., 1995. Functional connectivity in the motor cortex of resting human brain using echoplanar MRI. Magn. Reson. Med. 34, 537 – 541

34. Fox, M. D., & Raichle, M. E. (2007). Spontaneous fluctuations in brain activity observed with functional magnetic resonance imaging. Nat. Rev. Neurosci., 8, 700–11

35. Smith, S. M. et al. Advances in functional and structural MR image analysis and implementation as FSL. Neuroimage 23, S208–S219 (2004).

36. Marek, S., & Dosenbach, N. The frontoparietal network: function, electrophysiology, and importance of individual precision mapping. Dialogues in clinical neuroscience, 20(2), 133–140 (2018).

37. Vossel, S., Geng, J. J., & Fink, G. R. Dorsal and ventral attention systems: distinct neural circuits but collaborative roles. The Neuroscientist: a review journal bringing neurobiology, neurology and psychiatry, 20(2), 150–159 (2014).

