## Supplemental Tables for "The functional connectivity of the human claustrum according to the Human Connectome Project database"

**Supplementary Material**

**TABLE S1**. Left claustrum resting-state functional connectivity in the entire sample (N=100)

| **Brain Areas** | **Cluster size**  **(voxels)** | **Peak MNI**  **coordinates (mm)**  **x y z** | **Peak**  **T value** |
| --- | --- | --- | --- |
| Claustrum l/PreCentral Gyrus (l r)  PostCentral Gyrus (l r)  Insular Cortex (l r)  Central Opercular Cortex (l r)  Anterior Cingulate Cortex  Parietal Operculum Cortex (l r)  Supramarginal Gyrus (l r)  Supplementary Motor Area (l r)  Planum Temporale (l r)  Putamen (l r)  Inferior Frontal Gyrus (l r)  Heschl's Gyrus  Middle Frontal Gyrus (l)  Lateral Occipital Cortex (l)  Superior Frontal Gyrus (l)  Thalamus (l)  Amygdala (l) | 16897 | -34 -10 +00 | 23.44 |
| Lingual Gyrus (l r)  Intracalcarine Cortex (l r)  Precuneous  Cuneal Cortex (r)  Occipital Pole (l r) | 736 | -16 -46 -04 | 7.92 |
| Occipital Fusiform Gyrus (l)  Temporal Occipital Fusiform Cortex (l) | 177 | -24 -76 -08 | 6.79 |
| Cuneal Cortex (l) | 108 | -10 -78 +28 | 6.87 |
| Lateral Occipital Cortex (l) | 96 | -22 -86 +26 | 5.93 |

The results were corrected at seed level correction p<0.05 using False Discovery Rate (FDR).

l = left, r = right

**Table S2**. Right claustrum resting-state functional connectivity in the entire sample (N=100)

| **Brain Areas** | **Cluster size**  **(voxels)** | **Peak MNI**  **coordinates (mm)**  **x y z** | **Peak T value** |
| --- | --- | --- | --- |
| Claustrum r/ PreCentral Gyrus (r)  PostCentral Gyrus (r l)  Insular Cortex (r)  Cental Opercular Cortex (r)  Supramarginal Gyrus (r)  Parietal Operculum Cortex (r)  Anterior Cingulate Cortex  Supplementary Motor Area (r l)  Inferior Frontal Gyrus (r)  PlanumTemporale (r)  Frontal Operculum (r)  Middle Frontal Gyrus (r)  Paracingulate Gyrus (r)  Heschl's Gyrus (r)  Superior Frontal Gyrus (r)  Putamen (r) | 8544 | +34 +04 -02 | 18.40 |
| Planum Polare (l)  Central Opercular Cortex (l)  Precentral Gyrus (l)  Postcentral Gyrus (l)  Parietal Operculum Cortex (l)  Insular Cortex (l)  Planum Temporale (l)  Supramarginal Gyrus (l)  Frontal Operculum Cortex (l)  Heschl's Gyrus (l) | 2173 | -40 -22 +00 | 9.53 |
| Precentral Gyrus (l)  Middle Frontal Gyrus (l) | 121 | -30 -08 +48 | 7.17 |
| Lingual Gyrus (r)  Precuneous  Intracalcarine Cortex (r)  Cuneal Cortex (r) | 247 | +20 -56 +04 | 7.04 |

The results were corrected at seed level correction p<0.05 using False Discovery Rate (FDR).

l = left, r = right

**Table S3.** Claustrum resting-state functional connectivity

(Cls l vs. Cls r, cortical areas as ROIs)

| **Brain Areas** | **Beta** | **T (99)** | **p-FDR** |
| --- | --- | --- | --- |
| Insular Cortex (l) | 0.20 | 10.62 | 0.000000 |
| Frontal Orbital Cortex (l) | 0.12 | 8.37 | 0.000000 |
| Heschl's Gyrus (l) | 0.10 | 7.06 | 0.000000 |
| Frontal Operculum Cortex (l) | 0.10 | 6.69 | 0.000000 |
| Central Opercular Cortex (l) | 0.10 | 6.42 | 0.000000 |
| Inferior Frontal Gyrus (tri) (l) | 0.10 | 6.41 | 0.000000 |
| Frontal Pole (l) | 0.08 | 6.32 | 0.000000 |
| Paracingulate Gyrus (l) | 0.08 | 6.15 | 0.000000 |
| Middle Temporal Gyrus (to) (l) | 0.08 | 5.91 | 0.000000 |
| Parietal Operculum Cortex (l) | 0.09 | 5.75 | 0.000001 |
| Inferior Frontal Gyrus (oper) (l) | 0.09 | 5.67 | 0.000001 |
| Temporal Pole (l) | 0.10 | 5.66 | 0.000001 |
| Angular Gyrus (l) | 0.07 | 5.46 | 0.000002 |
| Planum Temporale (l) | 0.08 | 5.32 | 0.000004 |
| Middle Frontal Gyrus (l) | 0.07 | 5.23 | 0.000005 |
| Temporal Fusiform Cortex (a) (l) | 0.08 | 5.02 | 0.000012 |
| Supramarginal Gyrus (p) (l) | 0.07 | 4.99 | 0.000013 |
| Planum Polare (l) | 0.08 | 4.8 | 0.000027 |
| Middle Temporal Gyrus (a) (l) | 0.07 | 4.76 | 0.000030 |
| Superior Temporal Gyrus (a) (l) | 0.07 | 4.72 | 0.000034 |
| Superior Frontal Gyrus (l) | 0.07 | 4.60 | 0.000050 |
| Middle Temporal Gyrus (p) (l) | 0.06 | 4.60 | 0.000050 |
| Precentral Gyrus (l) | 0.07 | 4.50 | 0.000071 |
| Supramarginal Gyrus (a) (l) | 0.06 | 4.05 | 0.000357 |
| Parahippocampal Gyrus (a) (l) | 0.06 | 3.70 | 0.001211 |
| Superior Temporal Gyrus (p) (l) | 0.06 | 3.61 | 0.001555 |
| Postcentral Gyrus (l) | 0.06 | 3.54 | 0.001791 |
| Insular Cortex (r) | – 0.15 | – 8.93 | 0.000000 |
| Frontal Pole (r) | – 0.07 | – 4.24 | 0.000186 |
| Lateral Occipital Cortex (s) (r) | – 0.06 | – 3.60 | 0.001555 |
| Middle Frontal Gyrus (r) | – 0.05 | – 3.56 | 0.001718 |
| Superior Parietal Lobule (r) | – 0.05 | – 3.18 | 0.005557 |
| Inferior Temporal Gyrus (to) (r) | – 0.05 | – 3.14 | 0.006073 |

The results were corrected at seed level correction p<0.05 using False Discovery Rate (FDR).

l = left

r = right

a = anterior division

p= posterior division

s = superior division

tri= pars triangularis

to = temporooccipital part

oper = pars opercularis

**Table S4.** Claustrum resting-state functional connectivity (Cls l vs. Cls r, networks nodes as ROIs)

| Network | Brain Areas | **Beta** | **T (99)** | **p-FDR** |
| --- | --- | --- | --- | --- |
| Language | Superior Temporal Gyrus (p) (l) | 0.10 | 7.66 | 0.000000 |
| Language | Inferior Frontal Gyrus (l) | 0.10 | 6.83 | 0.000000 |
| Salience | A Insula (l) | 0.09 | 5.62 | 0.000002 |
| Salience | Supramarginal Gyrus (l) | 0.07 | 5.20 | 0.000009 |
| SensorioMotor | Lateral (l) | 0.07 | 4.31 | 0.000248 |
| Dorsal Attention | IntraParietal Sulcus (l) | 0.06 | 3.59 | 0.002322 |
| Dorsal Attention | Frontal Eye Field (l) | 0.04 | 3.12 | 0.008518 |
| Default Mode | Lateral Parietal (l) | 0.04 | 2.87 | 0.015968 |
| Salience | Rostral Prefrontal Cortex (l) | 0.04 | 2.61 | 0.028061 |
| Fronto-Parietal | Lateral Prefrontal Cortex (l) | 0.04 | 2.46 | 0.038658 |
| Fronto-Parietal | Lateral Prefrontal cortex (r) | – 0.06 | – 4.03 | 0.000589 |
| Dorsal Attention | Intraparietal Sulcus (r) | – 0.04 | – 2.78 | 0.019125 |

The results were corrected at seed level correction p<0.05 using False Discovery Rate (FDR).

**Table S5.** Claustrum resting-state functional connectivity (Cls l > Cls r, subcortical areas as ROIs)

| **Brain Areas** | **Beta** | **T (99)** | **p-FDR** |
| --- | --- | --- | --- |
| Putamen (l) | 0.17 | 9.29 | 0.000000 |
| Pallidum (l) | 0.08 | 4.73 | 0.000056 |
| Amygdala (l) | 0.08 | 4.28 | 0.000215 |
| Hippocampus (l) | 0.06 | 3.69 | 0.001368 |
| Thalamus (l) | 0.05 | 3.15 | 0.006453 |
| Hippocampus (r) | – 0.04 | – 2.64 | 0.023932 |
| Amygdala (r) | – 0.04 | – 2.51 | 0.026577 |

The results were corrected at seed level correction p<0.05 using False Discovery Rate (FDR).

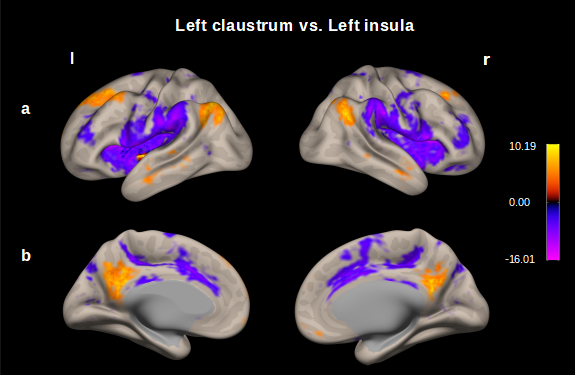

**Figure S1.** **Map of the resting-state functional connectivity left claustrum vs. left insula.** Seed – voxel analysis thresholded at p < 0.001 followed by p<0.05, p-FDR correction; entire sample 100 subjects.

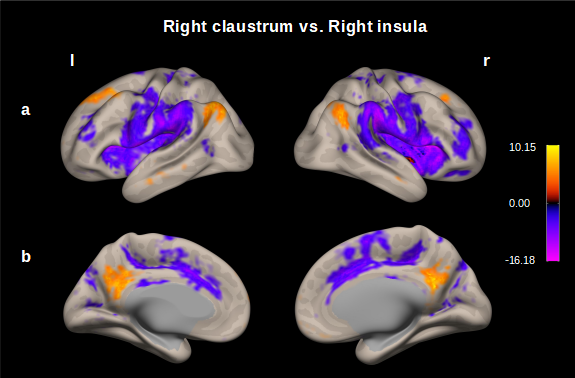

**Figure S2.** **Map of the resting-state functional connectivity right claustrum vs. right insula.** Seed – voxel analysis thresholded at p < 0.001 followed by p<0.05, p-FDR correction; entire sample 100 subjects.
